# An integrative genomic and chemical similarity approach linking fungal secondary metabolites and biosynthetic gene clusters

**DOI:** 10.1101/2025.05.08.652894

**Authors:** Karin Steffen, Manuel Rangel-Grimaldo, Thomas J. C. Sauters, David C. Rinker, Huzefa A. Raja, Tyler N. Graf, Adiyantara Gumilang, Olivia L. Riedling, Gustavo H. Goldman, Nicholas H. Oberlies, Antonis Rokas

## Abstract

Fungi are well known to biosynthesize structurally complex secondary metabolites (SMs) with diverse bioactivities. These fungal SMs are frequently produced by biosynthetic gene clusters (BGCs). Linking SMs to their BGCs is key to understanding their chemical and biological functions. Reasoning that structural similarity of SMs arises from similarities in the genes involved in their biosynthesis, we developed an integrative approach that leverages known BGC–SM pairs to predict global links across SMs and BGCs in fungi. As proof of concept, we systematically interrogated metabolomes and genomes of 16 strains of the filamentous fungus *Aspergillus fischeri*, detecting a total of 60 metabolites. Of those, we were able to assign 22 to known BGC–SM pairs and propose specific hypotheses for the remaining 38 metabolites. These results suggest that coupling genomic similarity and chemical structure-based similarity is a straightforward and high-throughput approach for linking fungal SMs to their BGCs.

## Introduction

Secondary metabolites (SMs), also known as specialized metabolites, allelochemicals, effectors, extrolites, or natural products, are instrumental to fungal ecology^1^. Fungal SMs contribute to various functions, including micronutrient acquisition (e.g., siderophores such as ferrichrome^2^), defense (e.g., antibacterials such as penicillin^3^), and pathogenicity (e.g., virulence factors such as gliotoxin^4^ and ToxA^5,6^). By virtue of their potent bioactivities, SMs are essential to drug discovery pipelines^7,8^, and thus for medical and agricultural research more broadly^9^.

In fungal genomes, the pathways involved in SM biosynthesis typically contain a set of neighboring, co-regulated genes, collectively referred to as biosynthetic gene clusters or BGCs^1^. A typical BGC contains genes coding for ‘core’ or ‘backbone’ enzymes responsible for the biosynthesis of the scaffold of the SM, tailoring enzymes that modify the scaffold, and cluster-specific transcription factors and transporters^1^. The clustering and content of genes in fungal secondary metabolite pathways led to the development of many different methods to predict BGCs^10^. These include CASSIS, a tool for predicting BGCs around a given anchor (or backbone) gene^11^; CLOCI, which predicts BGCs based on co-occurring loci and orthologous clusters^12^; DeepBGC, a machine learning-based tool trained on BGC presence in prokaryotic genomes^13^; and protein domain-based tools like the popular antiSMASH^14^ that predict BGC presence using profile hidden Markov models targeting required biosynthetic domains, along with BGC class-specific rules^14,15^.

Widespread access of column chromatography coupled with mass spectrometry (i.e., LC-MS and LC-MS/MS or LC-MS^n^) has driven the annotation of metabolites within natural product extracts and even *in situ*^16–18^. Yet, due to technical limitations, the degree of certainty of an observation of a SM can vary based on the approach used^19^. Assigning the chemical identity, and hence structure, of compounds within an extract of a natural product can be categorized into four levels of certainty^20^: 1) identified compounds for which there are orthogonal supporting structural data, 2) putatively annotated for which there are matches to spectral libraries, 3) putatively characterized compound classes for which there are matches to the class of compounds, if not the specific compound, and 4) unknown compounds. For the purposes of this report, we focused on the identified compounds (#1 in the list above), where the compounds were isolated and characterized by mass spectrometry and NMR spectroscopy or there were matches to a dereplication database that was built upon fully characterized compounds^21,22^.

To date, more than 30,000 fungal metabolites have been characterized^23^, and genomic examinations suggest that there are likely millions of predicted BGCs in fungal genomes^24–27^. In contrast, there are only about 608 experimentally verified BGC–SM pairs in fungi^26,28–30^. This discrepancy arises largely because BGC–SM pairings are typically established on a case-by-case basis, since confirmation of their pairing requires experimental validation^15,31,32^. Thus, the SMs biosynthesized by predicted BGCs in fungal genomes are typically not known to science, and as such most of these BGCs are considered “orphans”. Similarly, the biosynthetic pathways responsible for the vast majority of characterized fungal metabolites are also not known, hindering efforts to study their bioactivities.

The very small number of known BGC–SM pairs, coupled with the much larger numbers of known fungal metabolites and predicted orphan BGCs in fungal genomes, underscores the need for methods and strategies to predict BGC–SM pairs. To bridge this gap between chemotype and genotype, several general and specific methodologies have been developed to associate SMs and their cognate BGCs^33,15,34,35,32^. At the core of these general approaches typically lies the independent identification of BGCs via predictions from the genome, and structural identification of SMs via metabolomics, followed by an algorithm to predict connections.

Importantly, many of these algorithms take advantage of the MIBiG database^28,30^, a community effort cataloguing BGCs and their SMs, which includes information on the gene/protein sequences of the BGC with their known or putative functions, the organism the BGC–SM pair was identified in, and the resulting SM structures and bioactivities.

Strategies have sought to scale up BGC–SM prediction by integrating larger metabolomics data sets. For example, “correlation-based” approaches statistically associate BGC or gene cluster family (GCF)–SM pairs based on co-occurrence patterns^34^, while “feature-based” approaches rely on specific, searchable attributes (e.g., core enzymes, transcription factors or metabolomic spectral features like fragments and isotopes) to generate forward (BGC to SM) or reverse (SM to BGC) associations. These approaches have recently uncovered a novel class of BGCs, the isocyanide synthases^35^, and linked peptide natural products (e.g. ribosomally synthesized and post-translationally modified peptides (RiPPs) or non-ribosomal polyketide synthase (NRPS)) to their core genes^33,36,37^. Stable isotope labelling has also been used to connect mass spectrometric features (“compounds/SMs”) to BGCs by tracing known BGC substrates^38^.

Here, we introduce a novel strategy to link the chemical structures of experimentally identified SMs to their cognate BGCs by way of known SM-BGC relationships using structural similarity clustering. We then applied this strategy to the metabolomes and genomes of 16 strains of the filamentous fungus *Aspergillus fischeri* and the known BGC–SM pairs in the MIBiG database. This enabled us to confidently assign more than one third of detected metabolites to known BGCs that are present in *A. fischeri* genomes, and generate testable BGC-SM hypotheses in a straightforward, fast and *ab initio* manner for nearly all the remaining SMs. Our results suggest that coupling genomic similarity and chemical structure-based similarity is a powerful approach for linking of detected SMs to their BGCs in fungal genomes.

## Results

### Leveraging genomic and chemical similarity to infer BGC–SM pairs

To link SMs to their BGCs, we developed an integrative approach based on SM structural similarity (**Figure 1**). Specifically, we rely on two key components, the MIBiG database and SM structural clustering to generate hypotheses and link BGCs and SMs. Leveraging the MIBiG database allows us to connect BGC genes and SMs via their BGC accession ID^28^. Conceptually, structure similarity clustering is analogous to a BLAST search, in that it may result in an exact database match, as well as a range of increasingly dissimilar “hits”. In practice, structural similarity in small molecules is typically assessed via fingerprinting (i.e., generating a bit vector) of a structure, e.g., via SMILES (simplified molecular-input line-entry system, i.e. text abstractions of 2D or 3D structures of molecules)^27,39,40^. The similarity between a pair of fingerprints is then expressed using the Tanimoto (Jaccard) index, which is the ratio of the number of shared fingerprint bits to the union of bits in a pairwise comparison. This chemical structure similarity-based approach allows us to generate testable hypotheses in a high-throughput manner, without the need for large data sets, advanced computational infrastructure, additional knowledge, or specific experimental setups.

**Figure 1:**
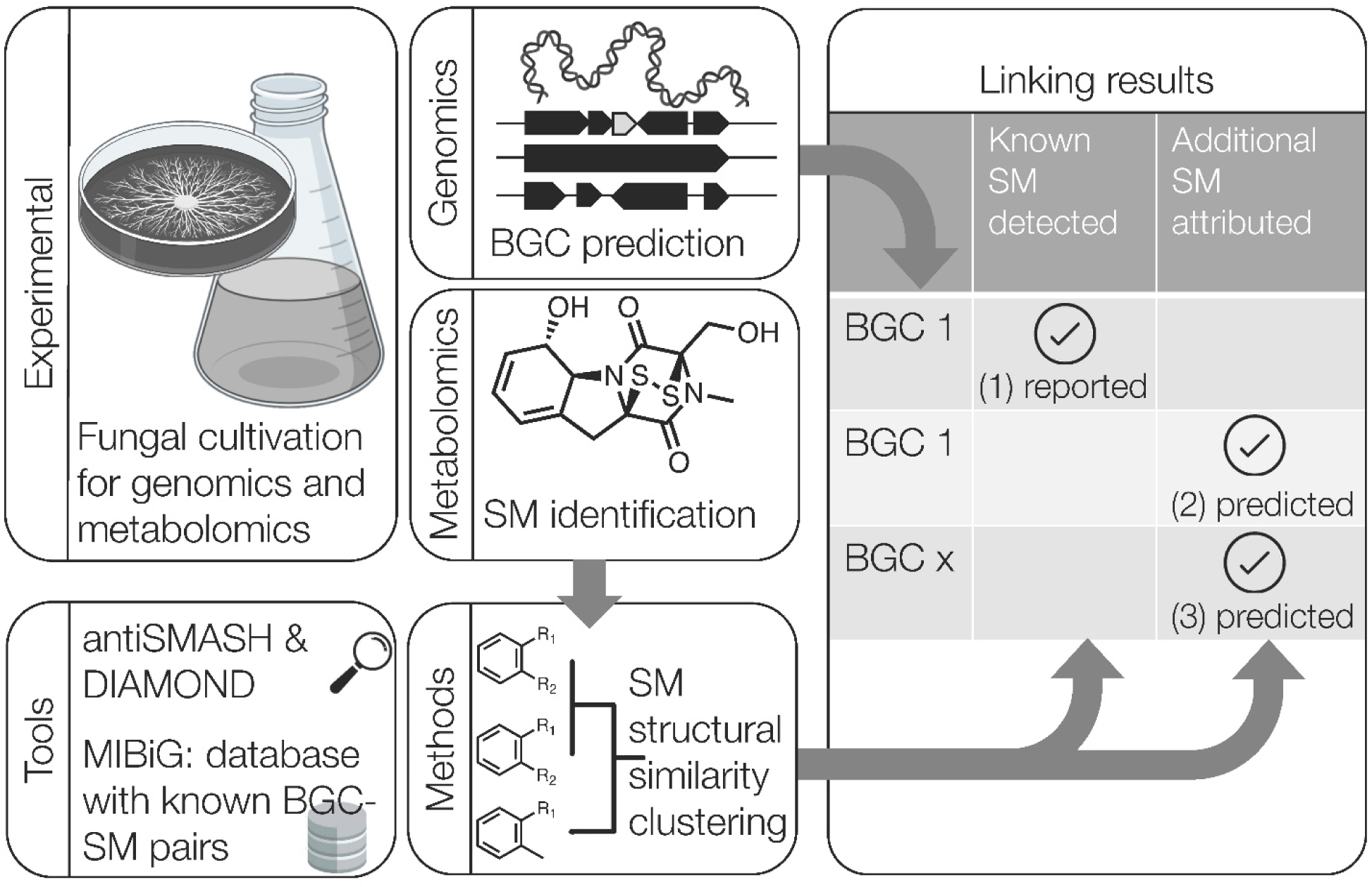
Schematic of workflow and components of the BGC–SM co-analysis. Fungal strains were cultivated for genomic and metabolomic analyses including detection of SMs and predicting BGCs, which were ultimately combined. Structures of experimentally identified SMs were clustered with those of known biosynthetic origin (from the MIBiG database), to (1) enable recovery of known BGC–SM pairs, (2) generate hypotheses about additional BGC–SM pairs, and (3) suggest groups of structurally similar SMs (SM groups) that are likely encoded by novel BGCs (i.e., not present in the MIBiG database). This pairing approach also includes orthogonal validation of *in silico*-predicted BGCs providing a focused and reliable view of biosynthetic capacities of the fungi.

### Sixty structurally characterized metabolites from *A. fischeri*

To test our method, we systematically interrogated the metabolomes and genomes of 16 strains of *Aspergillus fischeri*, a filamentous fungus that produces a variety of SMs and is gaining attention as a close, non-pathogenic relative of the major human pathogen *Aspergillus fumigatus*^41–43^. We assessed SM production at two temperatures (30°C and 37°C) using UPLC-MS/MS, as reported recently^43^. Compounds were identified based on either a direct match in LC-MS/MS to reference standards, all of which had been fully characterized by NMR, or to a class of fungal metabolites (i.e., via mass defect filtering). This experiment resulted in the positive identification of 60 compounds at two levels of confidence (**Table 1**, ‘A’ and ‘B’ respectively). Three biological replicates provided insight into the consistency of SM production by the various BGCs and strains (**Figure S1**). Overall, we found the most biosynthetically rich strains across all replicates and temperatures yield up to three times more SMs than the least-producing strains (CBS 150748: N=45, CBS 150751: N=42, CBS 150753: N=37 vs. CBS 147335: N=18, CBS 150754: N=15, CBS 54465: N=15). The compounds most consistently detected were helvolic acid and 16-*O*-deacetyl helvolic acid 21,16-lactone at both temperatures, azonapyrone A at 30°C, and acetylaszonalenin and aszonalenin at either temperature. Interestingly, strains with the greatest consistency of SM production across all biological replicates produced fewer compounds: CBS 150750 at 30°C (90%, 18/20 SMs were detected in all replicates), CBS 150757 at 37°C (85.7%, 18/21 SMs detected across all replicates), and CBS 150749 at 30°C (81.0%, 17/21 SMs detected across all replicates) (**Figure S2**).

**Table 1:** The 60 compounds identified from 16 strains of *A. fischeri* were grouped into 25 SM groups based on structural similarity. Each group was assigned an arbitrary identifier (i.e., 1 to 25). The superscript after the SM name indicates the level of experimental support: **^A^** MS/MS and NMR or MS/MS and dereplication with in-house database/standard; **^B^** MS/MS only. For each SM, the BGC(s) linked via compound clustering are indicated, with the underlined BGCs indicating the BGC of the SM resulting in the highest Tanimoto similarity. Pairings that could be validated based on experimental data (e.g., identical SM structures, evidence from the literature) are denoted as ‘reported’, and all newly generated hypotheses are denoted as ‘predicted’. For SMs of known BGCs, all generated hypotheses were accurate. For an overview of all structurally similar metabolites from *A. fischeri* together with their top SM hits in MIBiG database, where available, see **Figure S5**.

| SM group # | SM | BGC link | BGC present | confidence | Reference |
| --- | --- | --- | --- | --- | --- |
| 1 | Ilicicolin E <sup>B</sup> | <u>BGC0001923</u>,<br>BGC0001924 (New BGC 1) | no | predicted | 46 |
| 2 | (3β,22E)-Ergosta-4,6,8(14),22-tetraene-3-ol <sup>A</sup> | (primary metabolism) | – | – | 44 |
| 3 | Fumagillol <sup>B</sup> | BGC0001067 | yes | reported | 47 |
| 4 | Brevianamide A/B <sup>B</sup> | <u>BGC0001084</u>,<br>BGC0000816 (New BGC 2) | no | predicted |  |
| 4 | Brevianamide C/D <sup>B</sup> | <u>BGC0001084</u>,<br>BGC0000816 (New BGC 2) | no | predicted |  |
| 5 | Brevianamide Q <sup>B</sup> | BGC0000442 (New BGC 3) | no | predicted |  |
| 5 | Brevianamide R <sup>B</sup> | BGC0000442 (New BGC 3) | no | predicted |  |
| 5 | Brevianamide T <sup>B</sup> | BGC0000442 (New BGC 3) | no | predicted |  |
| 5 | Brevianamide U <sup>B</sup> | BGC0000442 (New BGC 3) | no | predicted |  |
| 5 | Brevianamide V/W <sup>B</sup> | BGC0000356 (New BGC 3) | no | predicted |  |
| 5 | Brevianamide K <sup>B</sup> | BGC0000442 (New BGC 3) | no | predicted |  |
| 6 | Cottoquinazoline E <sup>A</sup> | BGC0000355 | putative | predicted | 48,49 |
| 6 | Cottoquinazoline F <sup>A</sup> | BGC0000355 | putative | predicted | 48,49 |
| 6 | Cottoquinazoline G <sup>A</sup> | BGC0000355 | putative | predicted | 48,49 |
| 7 | Fumitremorgin F <sup>B</sup> | BGC0001142,<br>BGC0000355 (New BGC 4) | no | predicted | 50 |
| 7 | Fumitremorgin G/L <sup>B</sup> | BGC0001142,<br>BGC0000355 (New BGC 4) | no | predicted | 50 |
| 8 | 4-Hydroxyaszonalenin <sup>B</sup> | BGC0000293,<br>(BGC0002272) | yes | predicted | 51 |
| 8 | Acetylaszonalenin <sup>A</sup> | BGC0000293,<br>(BGC0002272) | yes | reported | 51 |
| 8 | Aszonalenin <sup>A</sup> | BGC0000293,<br>(BGC0002272) | yes | reported | 51,52 |
| 9 | Isoroquefortine C <sup>B</sup> | BGC0000420 | yes | reported | 53,54 |
| 9 | Roquefortine C <sup>B</sup> | BGC0000420 | yes | reported | 53,54 |
| 10 | Brevianamide E <sup>B</sup> | BGC0002272,<br>BGC0002617 | no | predicted |  |
| 11 | 13-O-prenylfumitremorgin B <sup>A</sup> | BGC0000356 | yes | predicted | 55,56 |
| 11 | Brevianamide F <sup>B</sup> | BGC0000356 | yes | reported | 55,56 |
| 11 | Deoxybrevianamide E <sup>B</sup> | BGC0000356 | yes | predicted | 55,56 |
| 11 | Fumitremorgin A <sup>A</sup> | BGC0000356 | yes | reported | 57 |
| 11 | Fumitremorgin B <sup>A</sup> | BGC0000356 | yes | reported | 55,56 |
| 11 | Fumitremorgin C <sup>A</sup> | BGC0000356 | yes | reported | 55,56 |
| 11 | spiro[5H,10H-dipyrrolo-[1,2-a:1',2'-d]pyrazine-2-(3H),2'-[2H]-indole]-3',5,10(1'H)trione <sup>A</sup> | BGC0000356 | yes | predicted | 58 |
| 11 | Tryprostatin B <sup>B</sup> | BGC0000356 | yes | reported | 55,56 |
| 11 | Tryprostatin C/D <sup>B</sup> | BGC0000356 | yes | predicted | 55,56 |
| 11 | Verruculogen <sup>B</sup> | BGC0000356 | yes | reported | 55,56 |
| 12 | hexadehydroastechrome (monomer) <sup>B</sup> | BGC0000372 | yes | reported | 59 |
| 12 | Trihistatin <sup>A</sup> | BGC0000420,<br>BGC0000372 | yes | predicted | 59 |
| 13 | 16-O-deacetyl helvolic acid 21,16-lactone <sup>B</sup> | BGC0000686 | yes | predicted | 60 |
| 13 | Helvolic acid <sup>A</sup> | BGC0000686 | yes | reported | 60 |
| 14 | Pyripyropene F <sup>B</sup> | BGC0000129,<br>BGC0001068 | yes | predicted | 61 |
| 14 | Pyripyropene H <sup>B</sup> | BGC0000129,<br>BGC0001068 | yes | predicted | 61 |
| 14 | Pyripyropene I <sup>B</sup> | BGC0000129,<br>BGC0001068 | yes | predicted | 61 |
| 14 | Pyripyropene O <sup>B</sup> | BGC0000129,<br>BGC0001068 | yes | predicted | 61 |
| 15 | Azonapyrone A <sup>A</sup> | BGC0002604 | yes | predicted | 62 |
| 15 | Sartorypyrone A <sup>A</sup> | BGC0002604 | yes | reported | 62 |
| 16 | Circumdatin C <sup>A</sup> | BGC0000355,<br>BGC0001652,<br>BGC0000448,<br>BGC0000409,<br>BGC0000303 (New BGC 5) | no | predicted |  |
| 16 | Dimetoxycircumdatin C <sup>A</sup> | BGC0000355,<br>BGC0001652,<br>BGC0000448,<br>BGC0000409,<br>BGC0000303 (New BGC 5) | no | predicted |  |
| 17 | Betaenone E <sup>B</sup> | BGC0002165,<br>BGC0001264 (New BGC 6) | no | predicted | 63 |
| 17 | Betaenone G/I/J <sup>B</sup> | BGC0002165,<br>BGC0001264 (New BGC 6) | no | predicted | 63 |
| 17 | Betaenone H <sup>B</sup> | BGC0002165,<br>BGC0001264 (New BGC 6) | no | predicted | 63 |
| 18 | Clavaric acid <sup>B</sup> | BGC0001248 | yes | reported | 64,65 |
| 19 | Chaetoglobosin 542 <sup>B</sup> | BGC0002539,<br>BGC0000968,<br>BGC0001182 | yes | predicted | 66 |
| 20 | Neosartoricin <sup>B</sup> | BGC0001144 | yes | reported | 67,68 |
| 20 | Neosartoricin C <sup>B</sup> | BGC0001144 | yes | reported | 67,68 |
| 20 | Neosartoricin D <sup>B</sup> | BGC0001144 | yes | reported | 67,68 |
| 21 | Brevianamide L <sup>B</sup> | BGC0002208,<br>BGC0002242 (New BGC 7) | no | predicted | 69 |
| 21 | Brevianamide O <sup>B</sup> | BGC0002208,<br>BGC0002242 (New BGC 7) | no | predicted | 69 |
| 21 | Brevianamide P <sup>B</sup> | BGC0002208,<br>BGC0002242 (New BGC 7) | no | predicted | 69 |
| 22 | Secalonic acids (A/ B/ C/ D/ E/ F/ F1/ G; 4,4'-Secalonic acid E) <sup>B</sup> | BGC0002063,<br>BGC0001886,<br>BGC0001988 | yes | reported | 70 |
| 23 | Nidiascin C <sup>A</sup> | BGC0002275,<br>BGC0002171 (New BGC 8) | no | predicted |  |
| 24 | Neosartorin <sup>A</sup> | BGC0001988 | yes | reported | 71 |
| 25 | Bisdethiobis(methylthio)-gliotoxin <sup>A</sup> | BGC0000361 | yes | reported | 4 |
| 25 | Gliotoxin <sup>A</sup> | BGC0000361 | yes | reported | 4 |

### SM structural clustering with reference data is a fast and straightforward method for linking SMs

To generate hypotheses about the biosynthetic origin of SMs from *A. fischeri*, we clustered their structures with all SMs in MIBiG based on structural similarity. All 60 experimentally identified chemical structures from *A. fischeri* were analyzed alongside known SMs from the MIBiG database, which contains 3,158 structures from 1,896 bacterial and eukaryotic BGCs, including 692 SMs from 377 BGCs from fungi. Morgan fingerprinting, pairwise structure similarity calculation using the Tanimoto index, and hierarchical clustering were then applied to place these 60 metabolites from *A. fischeri* in a context of known SMs (and their BGCs) from MIBiG, subsequently referred to as SM groups (**Figure 2; Figure S3**).

**Figure 2:**
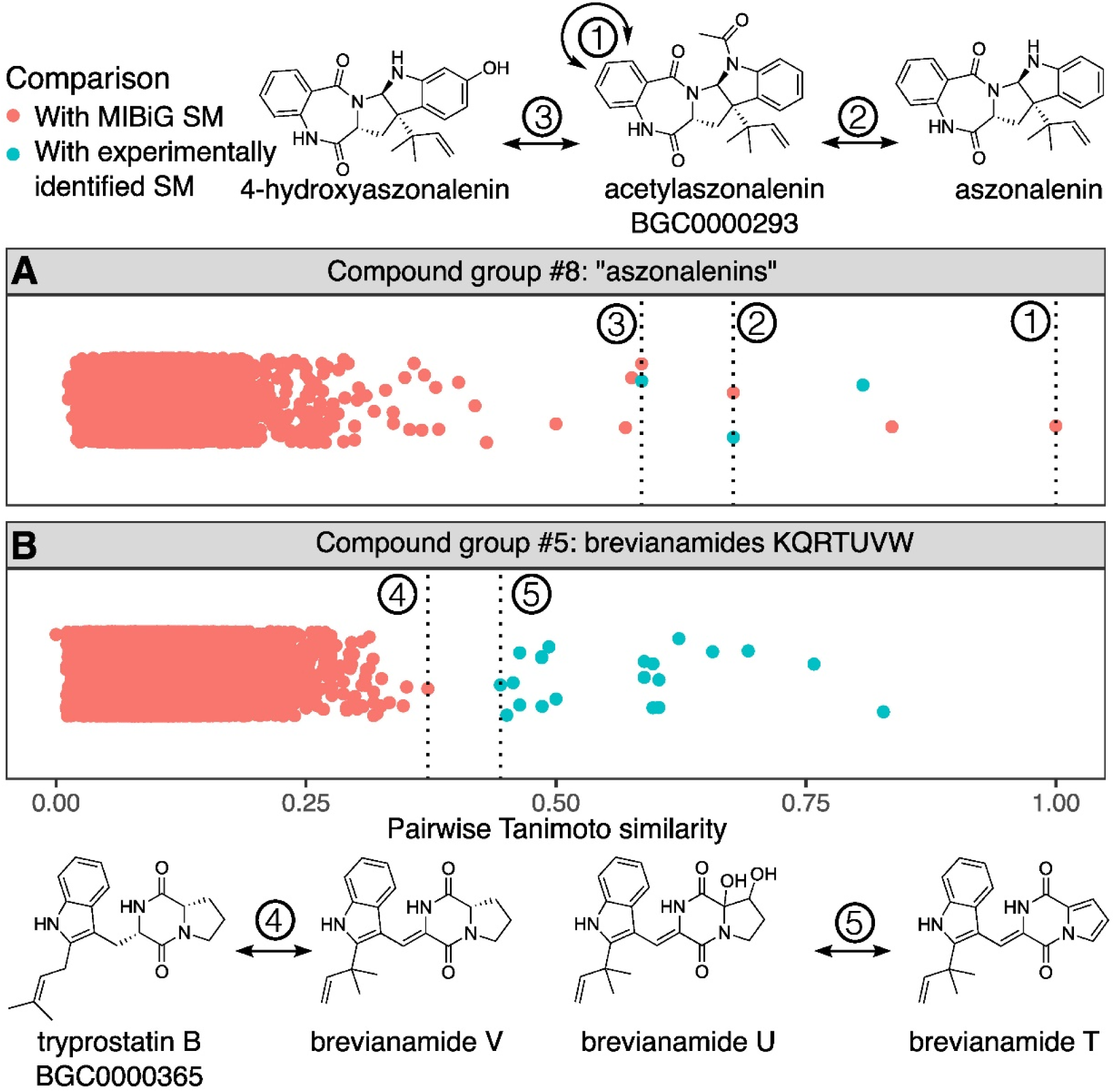
Evaluating SM groups by investigating pairwise similarities between secondary metabolites (SMs) within an SM group (blue) and with all SMs in MIBiG (red). Each dot represents the pairwise similarity between two SM structures, with higher values indicating greater similarity. **A) Matching an SM group to a known SM in the MIBiG database.** Three experimentally identified SMs, all aszonalenin analogs, have high pairwise similarities to each other (blue dots, compound group #8, Tanimoto similarities 0.59–0.81), which led us to group them. We show the similarity of each of the three compounds in this group to known MIBiG SMs as red dots, the dashed lines indicate the highest match between each experimentally identified SM and a MIBiG SM. As the identified SM acetylaszonalenin is also listed in MIBiG (BGC0000293), the comparison of two identical structures yields a similarity of 1 (denoted as 1). This SM yields also the highest similarity to aszonalenin (denoted as 2) and 4-hydroxyaszonalenin (denoted as 3). In summary, we define these three experimentally identified SMs as an SM group and link them to BGC0000293. **B) Proposing that an SM group represents a new SM (i.e., an SM absent from the MIBiG database).** Seven experimentally identified SMs, the brevianamides K, Q, R, T, U, V, and W yield high pairwise similarities amongst themselves (21 blue dots, compound group #5, Tanimoto similarities 0.44–0.83), and were thus placed into a single SM group. We show the pairwise structural similarities of all seven compounds with all MIBiG compounds as red dots. The highest similarity between any MIBiG SM and any of the experimentally identified SMs is tryprostatin B and brevianamide V (denoted as 4). This similarity is lower than the lowest similarity within the SM group (that between brevianamides U and T, denoted as 5). In summary, we define these seven experimentally identified SMs as group, but do not link them to any previously known SMs. While sharing the same diketopiperazine core made of L-Trp and L-Pro, all SMs in this group have distinct prenylations and other differences.

Based on identifying the most structurally similar SM in the MIBiG database for each identified *A. fischeri* metabolite, we hypothesize that their BGCs share similar gene content and organization. In doing so, we generated BGC hypotheses for all 60 identified metabolites from *A. fischeri*. Of these, 22 *A. fischeri* metabolites were identical to SMs in the database, i.e., representing known BGC–SM links, and 37 metabolites yielded biosynthetic hypotheses proposed based on structural similarity (**Table S1 ‘confidence’ column:** ‘reported’ and ‘predicted,’ respectively). The sole remaining metabolite is a sterol, which was not linked to a BGC, as sterol biosynthesis is part of primary metabolism^44,45^. The full hierarchical clustering analysis is presented in **Figure S3,** and background distributions of similarities of SMs produced by the same BGC and by different BGCs in **Figure S4**. **Figure 2** shows the pairwise similarities of SMs from two SM groups with all other SMs in MIBiG to illustrate: (panel A) a case of high similarity matches, which confirms the link between an experimentally detected *A. fischeri* metabolite and a known SM present in the MIBiG database, and (panel B) a case of low similarity matches, which suggests that the experimentally detected *A. fischeri* metabolites are likely biosynthesized by a new BGC, i.e. a BGC not currently present in the MIBiG database. The SM groups and hypotheses are described in **Table 1**, and are subsequently evaluated more generally.

### Assigning BGCs to identical pairs of structures

Thirteen SMs were identified from *A. fischeri* that had a corresponding SM structure included in the MIBiG database, and these formed SM groups together as expected. While unsurprising and seemingly trivial, the ability of our approach to quickly assign BGCs for experimentally identified SMs that are also present in the MIBiG database offers considerable practical utility, since the natural products literature does not dictate a consistent nomenclature process for SMs. For example, even compounds that could be considered analogues of each other are often named quite differently^72^, and this challenge is even greater for compounds that were discovered before the advent of computer-based searching (i.e., before SciFinder), where the same compound was often published under more than one name. A prominent example from the plant literature is salvinorin A (reported in 1982)^73^ and divinorin A (reported in 1984)^74^; these are the same compound, with the term ‘salvinorin’ now taking precedence^75^. However, both names can be found in the literature. Given that there are countless other examples, structure-based searches avoid identification issues stemming from ambiguous or differing nomenclature for both structurally identical and structurally similar compounds.

The 13 SMs identified from *A. fischeri* that had an identical SM match in the MIBiG database are: acetylaszonalenin, brevianamide F, clavaric acid, fumagilol, fumitremorgin B and C, helvolic acid, hexadehydroastechrom, neosartorin, roquefortine C, sartorypyrone A, tryprostatin B, and verruculogen. Notably, three known BGC–SM pairs were missed by our clustering approach due to database limitations. These were bisdethiobis(methylthio)gliotoxin and gliotoxin, both produced by BGC0000361, which was retired in MIBiG v3.1, and secalonic acid(s) for which the SM(s) were not structurally identified in our study nor in the existing MIBiG BGC0001886 entry.

### Uncovering BGCs for SMs not present in the MIBiG database

Not all known early biosynthetic intermediates, shunt products or possible SMs are deposited in the MIBiG database. Thus, when investigating BGC–SM hypotheses in SM groups, six additional BGC–SM links were confirmed based on primary literature. These are aszonalenin in BGC0000293^51^, fumitremorgin A in BGC0000356^57^, isoroquefortine C in BGC0000420^53^, and neosartoricin, neosartoricin C, and neosartoricin D in BGC0001144^67^. Notably, isoroquefortine C is an artifact produced by the isomerization of roquefortine C caused by pH or light^53^. Similarly, neosartoricin C and D might be artifacts related to the production of neosartoricin B^67^. Indeed, artifacts – compounds that were isolated but are likely changed in structure from the true SM, possibly due to extraction solvents or sample handling – are a well-known challenge in the natural products literature^76^. Finally, fumitremorgin A is technically not considered a product of the verruculogen BGC (BGC0000356), as the gene encoding the FtmPT3 protein responsible for converting verruculogen to fumitremorgin A is not part of the BGC^57^. However, this variation in the degree of clustering is not unusual, and numerous examples of such ‘satellite’ genes involved in SM biosynthesis, which reside outside the BGC, are known^77^. Given that fumitremorgin A is produced from verruculogen, an SM of this BGC, it is reasonable to include it in the set of SMs attributed to BGC0000356. In summary, for 22 of the 60 SM features (36%), our approach automatically annotated their respective BGCs. To confirm BGC presence predicted by known BGC–SM pairs and evaluate the hypotheses generated for the 38 remaining compounds without known BGCs (described below), we next examined the BGC content of the *A. fischeri* genomes.

### Genomic characterization of *A. fischeri* BGCs

To evaluate the BGC–SM pair predictions generated by our chemical clustering approach, we examined whether the BGCs of the pairs were present in the *A. fischeri* genomes and proteomes. To this end, we first analyzed the 16 genomes using antiSMASH v7, which predicts ‘BGC regions’ –i.e., continuous stretches in the genome containing a BGC and other genes (**Figure 3**). For traceability, we also identified and grouped homologous BGCs across the individual genomes. Across all 16 genomes, antiSMASH predicted 44 BGC regions that corresponded to 42 unique BGCs (BGC0001248 and BGC0002710 were each detected in two regions of the genomes), as well as 20 candidate BGC regions (‘unnamed’ putative BGCs). The mean number of BGCs per strain was 53.3 (range 51–56), a number consistent with previous reports^78^. Note that we refer to these predicted BGCs by the accession numbers of their reference BGCs in the MIBiG database.

**Figure 3:**
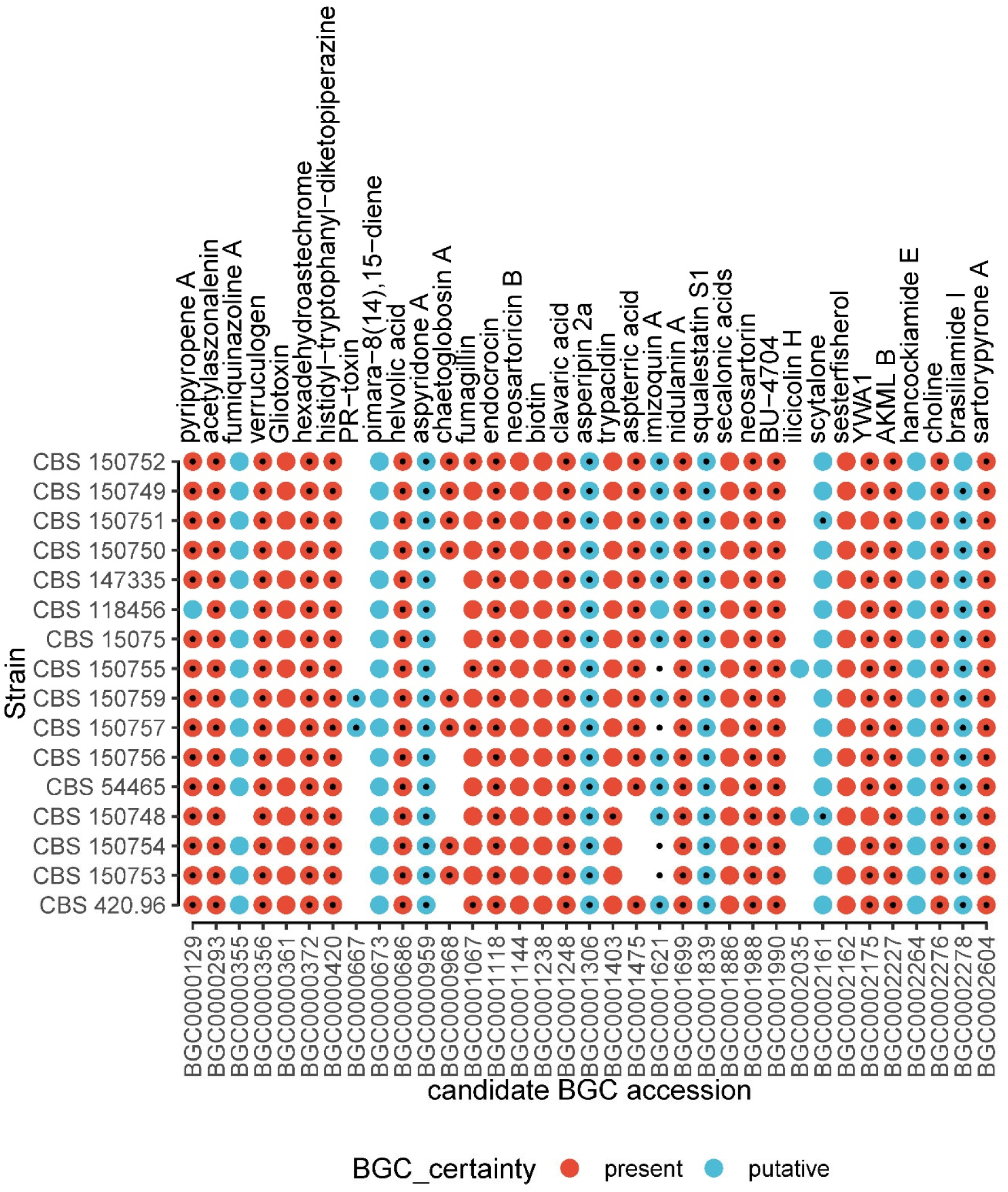
Map of BGC presence in *A. fischeri.* BGCs classified as ‘present’ (red) or ‘putative’ (blue) across 16 strains of *Aspergillus fischeri*. On top, a representative SM of each BGC is denoted for convenience. Large circles indicate the validated status (‘present’ or ‘putative’) of the BGC in the respective strain. Small black dots indicate BGCs detected by antiSMASH. The five artifactual BGCs, antiSMASH-predicted BGCs found ‘absent’, and unnamed BGC regions were not included. *A. fischeri* strains are ordered according to their phylogeny inferred by Rinker *et al*. ^43^. For a complete evaluation of all antiSMASH-predicted BGCs, see **Table S3**.

When comparing the correspondence of *A. fischeri* BGCs with their cognate reference sequences in the MIBiG database, BGCs were classified as ‘present’ in *A. fischeri* when they contained all the genes present in their MIBiG entry, or when they were incomplete but supported by evidence from chemistry studies (i.e., identified SMs). Additionally, BGCs were classified as ‘putative’ in *A. fischeri* when they were incomplete with at least half of the genes detected but without evidence from the chemistry studies. Otherwise, BGCs were classified as ‘absent’. Examining each of the 44 antiSMASH-predicted BGC regions across *A. fischeri* genomes, we classified 20 BGCs as ‘present’, 9 as ‘putative’, and 15 as ‘absent’ (**Table S3**). We also specifically searched for the protein sequences of each MIBiG BGC in the proteomes, allowing us to manually curate and validate the antiSMASH predictions. These additional analyses enabled us to classify 7 additional BGCs as ‘present’ and 4 BGCs as ‘putative’. A full list of BGCs is given in **Table S3**, with information on sequence identity with known BGC genes and genomic location in **Table S4**).

Among all 40 BGCs classified as ‘present’ or ‘putative’, five artifacts reduce the total BGC count. These primarily stem from the current cataloging approach for BGCs and our limited understanding of them. MIBiG defines each BGC in the genome it is reported in, sometimes listing the ‘same’ (i.e., homologous) BGC multiple times from different organisms. Similarly, one BGC that biosynthesizes one SM can be nested within another, larger BGC that biosynthesizes a different SM. These situations can lead to ‘collisions’, i.e. the assignment of the same set of genes to multiple BGCs.

There were collisions in two pairs of BGCs in our data where the same set of proteins in *A. fischeri* is classified as two different BGCs due to similarity of the MIBiG reference sequences (BGC0000361 gliotoxin / BGC0001609 gliovirin, and BGC0001144 neosartoricin B / BGC0002646 hancockinone A), reducing the unique BGC count by two. The BGC for biotin is listed twice in the MIBiG database (BGC0001238 and BGC0001239) but was counted only once, as it matches the same set of genes. Similarly, there are two slightly different BGCs matching a congruent set of genes for the compound ilicicolin H (BGC0002035 and BGC0002093), which were counted as one BGC, further decreasing the total count by two.

Additionally, the BGC for clavaric acid (BGC0001248), which is composed of a single gene, was found twice (**Table S4**). However, only one of the two homologs identified (homolog ID 221721_1) contains the sequence motif VSDCISE, which was previously found in *Fusarium graminearum* to be involved in clavaric acid production^65^.

In total, we infer that *A. fischeri* contains 35 ‘present’ and ‘putative’ BGCs (**Figure 3**, **Table S3**). Overall, BGC content was largely conserved and consistent across the 16 strains from diverse geographic locations^43^. The majority of BGCs, 82% (29/35 total ‘present’ and ‘putative’ BGCs) was detected in all strains with only 17% (6/35 BGCs) detected only in a subset of strains.

### Evaluating hypotheses for *A. fischeri* metabolites without identical matches to known SMs or mentions in the literature

We previously identified 22 metabolites with known BGCs. The remaining 38 experimentally detected *A. fischeri* metabolites did not have identical matches to SM structures included in the MIBiG database or biosynthetic information in the literature. Thus, we augmented the BGC–SM hypotheses based on structural similarity for each of these metabolites by examining whether *A. fischeri* genomes contained the SM-predicted BGCs (for details on BGC prediction/detection, see Methods). This evaluation resulted in predictions that we broadly grouped into three level-of-confidence categories: (i) attributing the metabolite to a known BGC that is present in the respective *A. fischeri* strain genome(s) (e.g. 4-hydroxyaszonalenin – BGC0000293, **Figure 2A**), (ii) linking the SM to a BGC not present in the respective *A. fischeri* genome(s). This implies either the presence of a homologous (but divergent) BGC or that the SM is produced by a convergently evolved, distinct BGC, and (iii) ascribing the SM (or SM group) as a novel metabolite(s) likely encoded by an unknown BGC (i.e., no similar SMs are present in the MIBiG database, **Figure 2B**). Given the dearth of fungal BGCs in MIBiG (i.e., only 377), we were pleased that our approach predicted 13 SMs in category (i), 11 SMs in category (ii), and 13 SMs in category (iii) (see extended **Table S1** for more details, and **Table S2** for all Tanimoto similarities). The one remaining metabolite, (3β,22E)-ergosta-4,6,8(14),22-tetraene-3-ol, is a sterol. Sterols are essential parts of eukaryotic membranes, and their biosynthesis cannot be linked to a BGC-SM pair since they are part of primary metabolism ^44^. Thus, cross-referencing the curated list of *A. fischeri* BGCs with the SM-predicted BGCs shows that 38 of the 59 BGC– SM links (64%) are orthogonally validated by the presence of the cognate or predicted BGC (**Table S1**).

### Caveats for using antiSMASH as tool for accurate BGC surveys

AntiSMASH is a widely used tool for BGC prediction in fungal genomes^32^. While examining the correspondence between *A. fischeri* BGCs identified by antiSMASH and their inferred references in the MIBiG database, we noted several idiosyncrasies that could lead to inaccurate predictions. In most known BGCs, the genomic ‘region’ predicted by antiSMASH to contain a BGC was much larger (up to approximately three times the number of genes) than the actual BGC. As BGCs are known to co-localize, particularly in telomeric or low complexity regions of genomes^79,80^, their physical proximity on chromosomes, in combination with the ‘regions’ concept of antiSMASH leads to BGCs masking each other (**Figure 4A, 4B**). This masking occurred in the proximal BGCs BGC0001403 for trypacidin and BGC0001988 for neosartorin, and with BGC0000356 for verruculogen and BGC0001067 for fumagillin. Another caveat when using antiSMASH with the module ‘--cb-knownclusters’ is the incompleteness of the MIBiG database used for the BGC prediction. Curation processes continually expand the knowledge base^30^, but sometimes, valid BGCs are removed or lacking, thus leading to missed predictions (e.g., the gliotoxin BGC, which was retired in MIBiG v3.1/v4.0). Examination of 16 strains of *A. fischeri* revealed some instances where the same homologous genes were predicted as part of different BGCs in different strains (**Figure 4C**). Finally, we noted that some BGCs were missed by antiSMASH (but detected by protein sequence searches), for no apparent reason (**Figure 3**).

**Figure 4.**
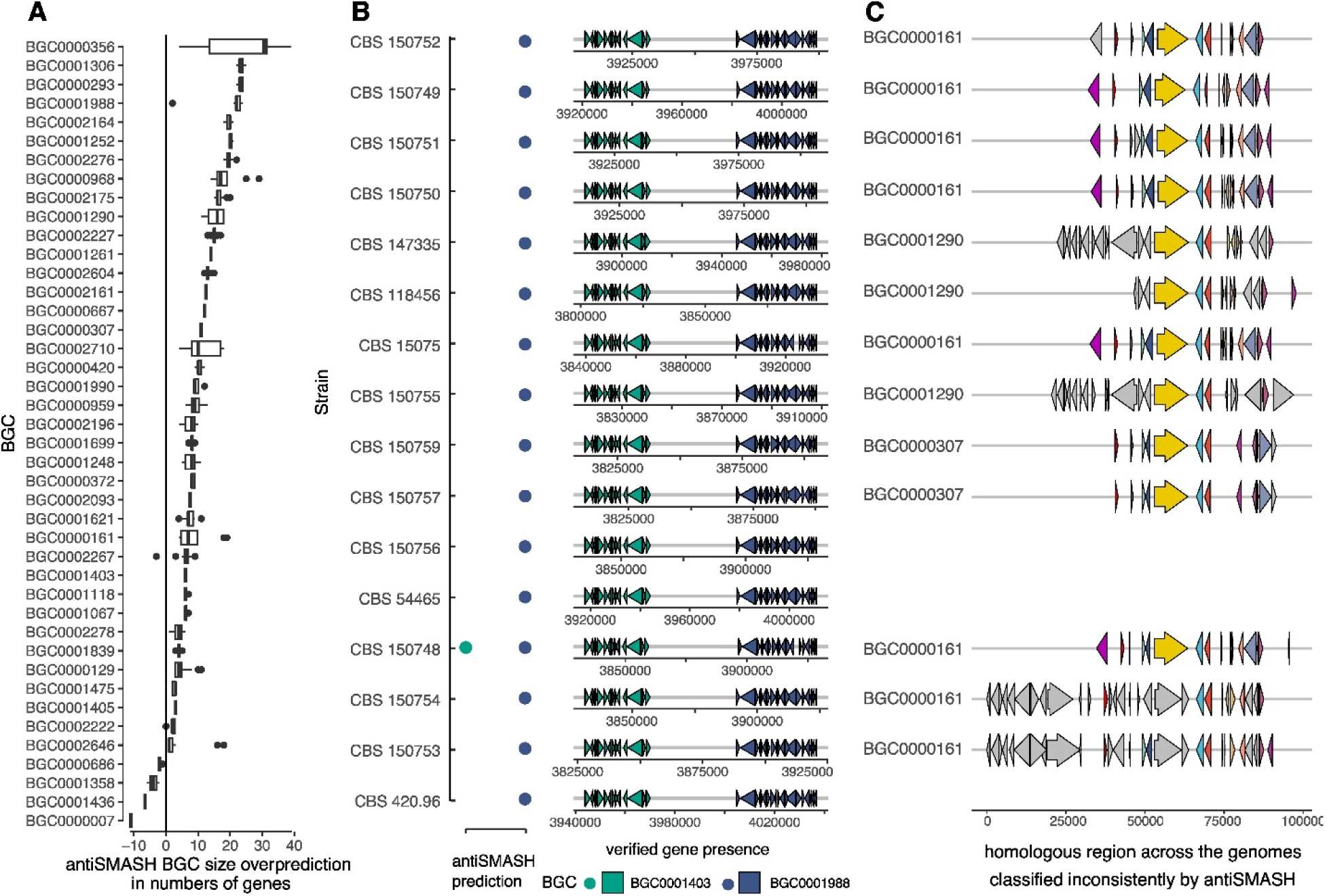
Inaccuracies in antiSMASH predictions of individual BGCs in fungal genomes. Properties leading to inaccurate BGC assessments include: (A) region overprediction, (B) merging/masking, and (C) inconsistent predictions. Panel **A**: BGC ‘region’ overprediction, calculated as the difference between the number of genes in the antiSMASH-predicted ‘region’ and the true number of genes in the BGC. These regions often include the BGC along with neighboring genes that contain putatively biosynthetic domains. Overprediction can lead to artifactual merging of neighboring BGCs (see panel B). Panel **B**: Example of artifactual merging and masking of BGC0001403 and BGC0001988. The two clusters are near each other in *A. fischeri* genomes (gene diagram), resulting in the merging of BGC0001988 into the prediction of BGC0001403 in all but one of the genomes (dots). We find both BGCs present in all strains, but antiSMASH missed 15 of 16 instances of BGC0001403. Panel **C**: Inconsistent assignments. The strains are the same as in panel B. The y-axis label indicates the BGC accession ID given by antiSMASH. Orthologous genes are indicated by color. Tracking orthologs across strains revealed that the same region was assigned to several different BGCs (BGC0000161, BGC0000307, BGC0001290). Here, all orthologs attributed to BGC0001290 were plotted, showing that a BGC region consisting of the same genes was predicted as three different BGCs across the strains.

## Discussion

Given the at least 30,000 reported fungal metabolites^23,81–83^ and millions of BGCs predicted in fungal genomes^25–27^ but the existence of only a few hundred BGC-SM pairs ^30^, linking BGCs^83^ and SMs is a persistent challenge.

In this work, we developed a chemical similarity approach that requires a minimum of input data. It can evaluate a single SM structure against the MIBiG database and querying of a single genome, or it can be scaled up as needed. The approach is independent of any specific experimental setup and can be performed entirely *in silico*, requiring only the SMILES structure of a compound of interest and a publicly available genome or proteome of the producing organism. Across 16 strains of a single fungal species, we recovered 22 known BGC–SM pairs and generated hypotheses for 37 more, including 11 additional SMs attributed to BGCs present. In our chemical similarity analyses, we refrained from setting a similarity threshold, due to the known patchiness of SMs present in the MIBiG database, general limitations in BGC–SM pairing knowledge, and previously documented challenges with threshold-based approaches^34^. Moreover, some SMs (particularly early biosynthetic intermediates) are not necessarily unique to any single BGC. As a result, these compounds can lead to different SM groups containing structurally similar SMs or identical similarity scores across different BGCs, thereby introducing ambiguity and ties if a strict threshold were applied. For example, the diketopiperazine brevianamide F (cyclo-L-Trp-L-Pro) is the first product of the biosynthesis of verruculogen by BGC000356 in *A. fumigatus*^55^ as well as of the biosynthesis of notoamide A by BGC0000818 in *Aspergillus versicolor*^84^ and brevianamide A by the *bvn* gene cluster in *Penicillium brevicompactum* NRRL 864^85^. The latter is currently not listed in MIBiG.

By contrast, existing approaches either require fairly large datasets or extensive experimental data. For example, using a metabologenomics approach, a recent correlation-based linking method generated hypotheses about BGC–SM pairs based on shared co-occurrence patterns identified via statistically significant correlations between 25 known SMs and 21 BGCs across 110 Ascomycete fungi^34^. While powerful, this approach requires fairly large datasets of BGCs and SMs. Similarly, the BGC-class agnostic strategy, IsoAnalyst^38^, predicts BGC–SM pairs by using stable isotope labels. This insightful approach cannot be applied for large-scale BGC–SM pairing, since it requires a specific experimental setup, where microbes are cultured for several days in replicates in the presence of the isotopically labelled substrates, along with knowledge of which substrate is utilized by each BGC^38^.

Additionally, the chemical similarity approach in this case study took less than 15 minutes on a laptop and is thus a very fast, accessible approach for generating testable biosynthetic hypotheses (BGC–SM pairs) in a high-throughput fashion. In cases where compounds (or groups of compounds) lack structurally similar SM neighbors whose biosynthetic pathways are known, our approach can suggest that they are likely the products of “novel BGCs”, facilitating efforts to discover new divergent genes and enzymes. Beyond the value of establishing BGC–SM links for understanding biosynthetic potential, this method lends orthogonal validation to BGC predictions, i.e., enabling prediction confirmation where known BGC–SM pairs are recovered. This increases the fidelity of the biological conclusions drawn based on the BGCs and their implications for the chemotype (i.e., SM profile), lifestyle, and niche of an organism.

When applying this chemical structure similarity approach in other contexts, there are a number of caveats. Our results are based on examination of strains of an *Aspergillus* species, which belongs to one of the most well studied fungal genera in terms of prior knowledge of BGC–SM pairs. Studies of less-studied species that are underrepresented in the MIBiG database may be more challenging, especially if their chemodiversity differs from the SMs currently represented in the MIBiG database. In general, the hypotheses (SM groups) may be a poor fit in the case of less well-studied species and compound classes in the database or when including other types of metabolites (e.g., primary metabolites like sterols). Additionally, chemical conversions that alter the backbone or skeleton of a SM sufficiently could mask a better clustering fit. Furthermore, when SMs are produced by multiple non-homologous BGCs (e.g., brevianamide F), genomic evidence is necessary to determine which BGC it is produced by. Such instances of convergently evolved SMs would only be detected in this strategy when finding the SM and not the BGC (but this inference would be based on the absence of evidence). Putative examples of this in our data are cheatoglobosin 542 and ilicicolin E. Cheatoglobosin 542 is structurally very similar to cheatoglobosin A produced by BGC0000968 ^86^. The *A. fischeri* BGC exhibits very high amino acid sequence similarity to this MIBiG BGC. However, the presence-absence patterns of the BGC and the SM do not match, which led us to a “second best” BGC, which has lower sequence similarity to the MIBiG BGC but is present in a SM-congruent pattern. In the second case, ilicicolin E differs from ascochlorin of BGC0001923^46^ only by the presence of an α,β-unsaturated ketone instead of the aliphatic ketone, respectively, in the 6-membered ring. However, there is no indication in the genomes of a similar or derived BGC. In both cases, SM structural similarity may result from convergent evolution rather than from genomic similarity.

These caveats notwithstanding, our approach successfully inferred BGC–SM pairs for nearly one third of the fungal metabolites identified and predicted BGC–SM pair hypotheses for nearly all the rest. Ultimately, our approach is a hypothesis-generating strategy and must be validated experimentally (e.g., by modifying putative BGCs in the native host or through heterologous expression of the putative BGC)^31,87^. The approach applied in this work leveraged similarity among known SM structures and BGCs to bidirectionally link SMs and BGCs via the MIBiG database and thereby successfully generated testable biosynthetic hypotheses in a high-throughput fashion.

## Methods

All genomic and metabolomic data were taken from Rinker *et al*.^43^

### Chemical fingerprinting and clustering

Structures (SMILES, simplified molecular-input line-entry system; a text string representing the molecule) for all SMs identified from untargeted metabolomics were collected via ChemDraw v23.1.1 (Revvity) and combined with structures from known BGCs deposited in the MIBiG database ^30^. Chemoinformatic analyses were carried out in Jupyter notebook^88^ using RDKit and PubChemPy^89^. To facilitate the subsequent search for the detected compounds, we prefixed the names of structures from MIBiG with the BGC accession ID, and those of compounds found in extracts with ‘chem_’. For comparing structural similarity and clustering the compounds, we calculated the Morgan fingerprint for each compound with GetMorganFingerprintAsBitVect() using chirality with a radius of 2, and 2048 bits, and converted the fingerprints to binary strings using ToBitString(). We calculated Tanimoto similarity (Jaccard index, the intersection of set bitflags divided by the union) between all pairwise comparisons using calculate_tanimoto() resulting in a symmetric similarity matrix of all-vs-all comparisons. With linkage(method=’average’, metric=’euclidean’) and dendrogram() from scipy, we performed hierarchical clustering of the compounds based on the distance matrix and used matplotlib to plot and save the resulting figure (**Figure S3**). SM groups were initially delineated by searching the dendrogram for the tag “chem_” and grouping similar structures. Subsequently, for every identified SM, the highest pairwise similarity scores with a SM in the MIBiG database was extracted from the similarity matrix, thereby generating biosynthetic hypotheses for each SM.

### BGC predictions

Genomic data were taken from Rinker et al.^43^ and BGCs were predicted using antiSMASH v7.1.0^14^ and DIAMOND v2.1.6.160 blastp searches^90^ of the MIBiG database v3.1^28^. All subsequent analyses were performed in R v4.4.0^91^. Conventionally, BGCs are defined in a specific genome. However, in this manuscript, we refer to the predicted candidate BGCs by their MIBiG accession number for convenience.

After the antiSMASH prediction (--fullhmmer--rre--cc-mibig--cb-knownclusters--cb-subclusters--cb-general, using the corresponding gff3 annotation file), an in-house script was used to aggregate results from individual runs. Across the different strains, known and unknown BGCs were aggregated by comparing gene content. This approach yielded meaningful clusters as evidenced by the correct grouping of known BGCs (with their MIBiG BGC accession ID). This clustering revealed instances in which the same genes were attributed to different BGCs (both known and “anonymous” candidate clusters) by antiSMASH.

For the amino acid sequence search, the 16 genomes were queried with all sequences in MIBiG v3.1 (diamond blastp -f6 qseqid sseqid pident length mismatch gapopen qstart qend sstart send evalue bitscore qcovhsp qlen slen full_sseq) and the results concatenated.

To validate the antiSMASH BGC predictions, the genes in each predicted region were searched using DIAMOND blastp. Additionally, the hits were filtered for high identity (pident >80%, minimum 50% query coverage), as well as for runs of hits against the same BGC in proximity (low identity clustering of putative, diverged BGCs). By using DIAMOND blastp to confirm *A. fischeri* BGC genes based on known BGCs, we tagged every BGC gene with a BGC ID from MIBiG hence allowing for interoperability of biological and chemical data.

BGCs were classified as present if all genes were found in proximity, regardless of whether a corresponding SM was detected, or if they were recovered partially, i.e. incomplete but with evidence from SMs. BGCs were classified as putative if more than half of the genes were present but there was no evidence for their presence based on metabolomics. BGCs were classified as absent if fewer than half of the genes were found and no evidence from metabolomics was present.

