## Supplementary Figures for "An integrative genomic and chemical similarity approach linking fungal secondary metabolites and biosynthetic gene clusters"

### Supplementary material

#### Readme

This document contains supplementary information for the publication "**An integrative genomic and chemical similarity approach linking fungal specialized metabolites and biosynthetic gene clusters**" by Steffen *et al.*

Under "**Supplementary information**", we compiled figures and tables mentioned in the main text and their legends. Exceptions are Figures S3 and S5 which are provided separately due to their size.

We further include information on supplementary tables (deposited as supplementary file with this manuscript), as well as raw and intermediate data files provided for reproduction (deposited in a Figshare repository due to their size). In the Figshare repository, you also find analysis code and a GUI, as outlined in "**Custom analysis code and in-house scripts**".

#### Supplementary Information

Integrating chemical evidence affords more nuanced insights into the phenotypes, and can be facilitated by encoding structural information via SMILES. In this study/In Rinker *et al.*, [1], the three biological replicates for metabolomics were treated as uniformly as possible. Despite best efforts, we find that the production of many SMs is somewhat inconsistent across the replicates and strains (**Figure S1**). This adds nuance to the notion of the 'silent BGC', as it seems that in *A. fischeri*, only a handful of BGCs are active consistently and detected across all three replicates (**Figure S2**). To detect a maximum diversity of SMs, it seems that biological replicates should be considered valuable and informative together with other established methods such as co-culturing [2,3] or cultivation on different substrates (OSMAC 'one strain, many compounds' [4]) to elicit a diverse chemical response by the fungus.

#### Supplementary figures

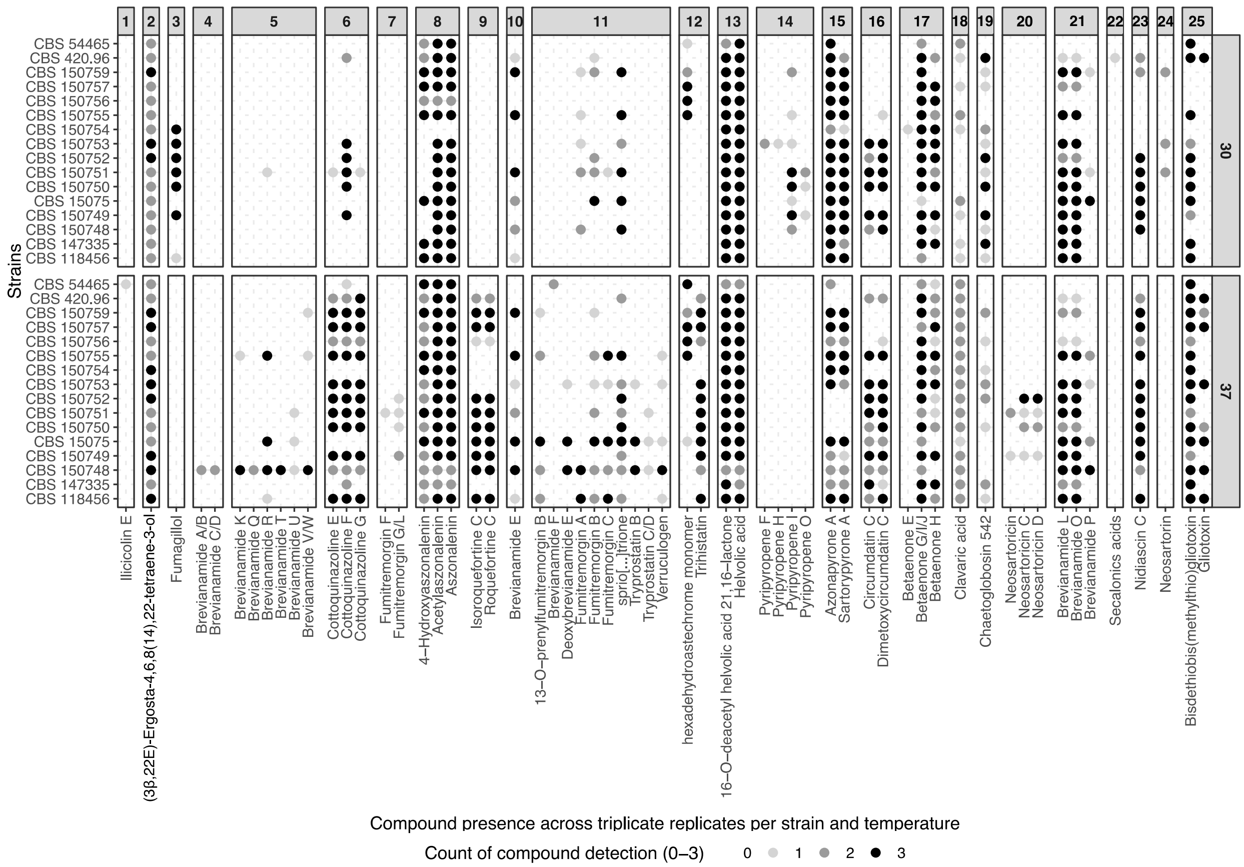

**Figure S1**: Per strain and temperature indication of SM expression/detection across the triplicate replicates. The SMs are grouped by assigned compound clusters. The compound ‘spiro[5H,10H-dipyrrolo-[1,2-a:1′,2′-d]pyrazine-2-(3H),2′-[2H]-indole]-3′,5,10(1′H)trione’ was abbreviated ‘sprio[...]trione’.

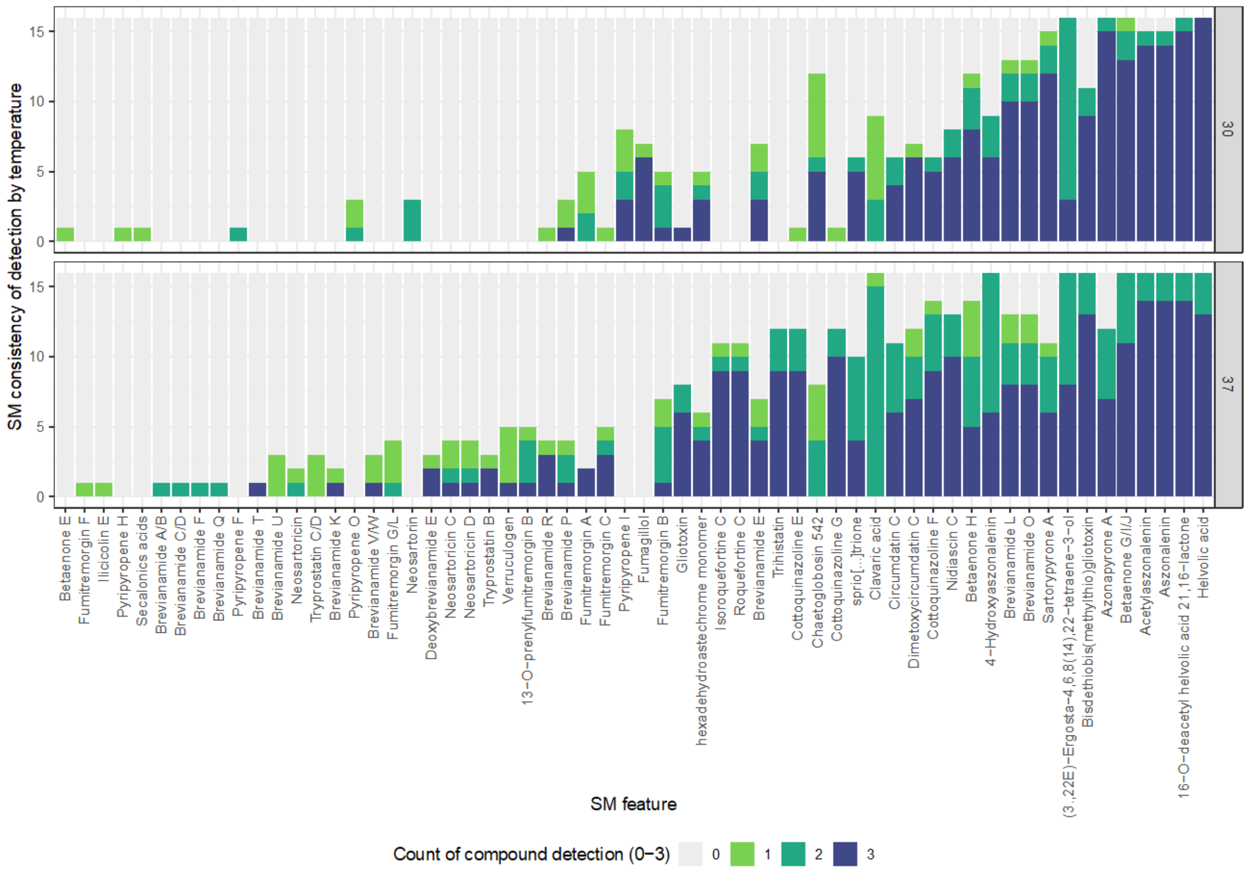

**Figure S2**: Consistency of expression/detection of SMs at 30°C and 37°C. Out of 16 strains, in how many strains was each SM detected consistently, i.e. in all replicates (3) or in none of the replicates (0), and in how many strains was the SM detected inconsistently (i.e. in one or two out of three replicates).

**Figure S3**: Dendrogram representing the hierarchical clustering of experimental SMs with all structures deposited in MIBiGv 3.1.

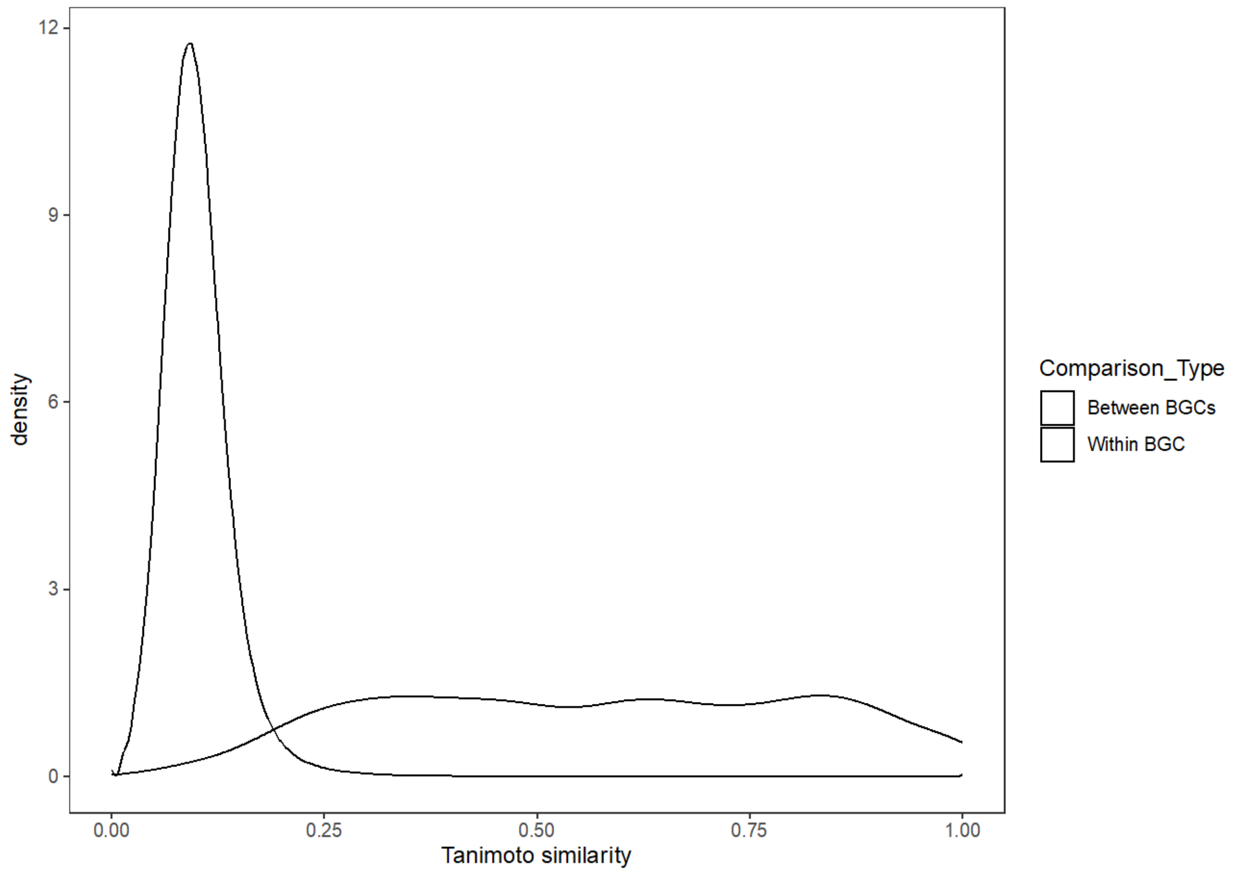

**Figure S4**: Background distribution of similarity scores for pairwise SM comparisons within a BGC (blue) and with compounds from other BGCs (red). For comparison, vertical lines indicate the similarity values of the best match for experimentally identified SMs with a MIBiG SM.

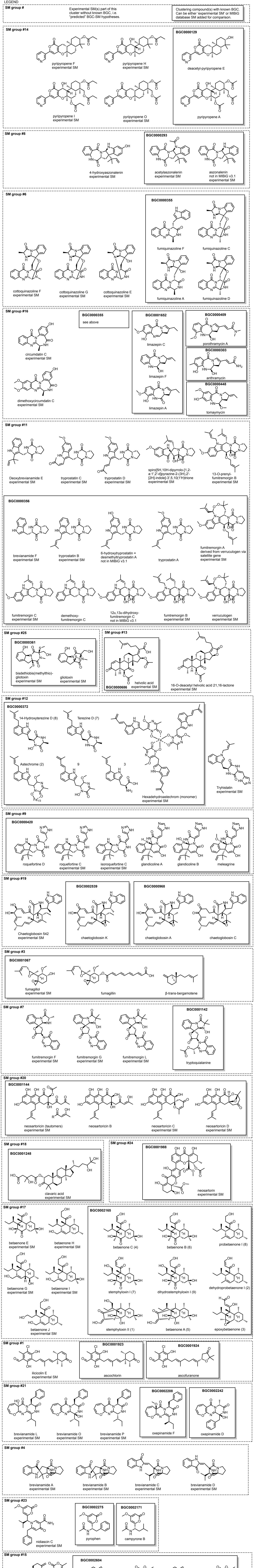

The multicyclic structure of Brevianamide E could be derived from Brevianamide W via a ring closure (between the indole and amide motifs) and subsequent hydroxylation. It could be speculated that this single bond difference affected the backbone and thus fingerprint enough to exclude brevianamide E from the cluster that it might biosynthetically belong to.

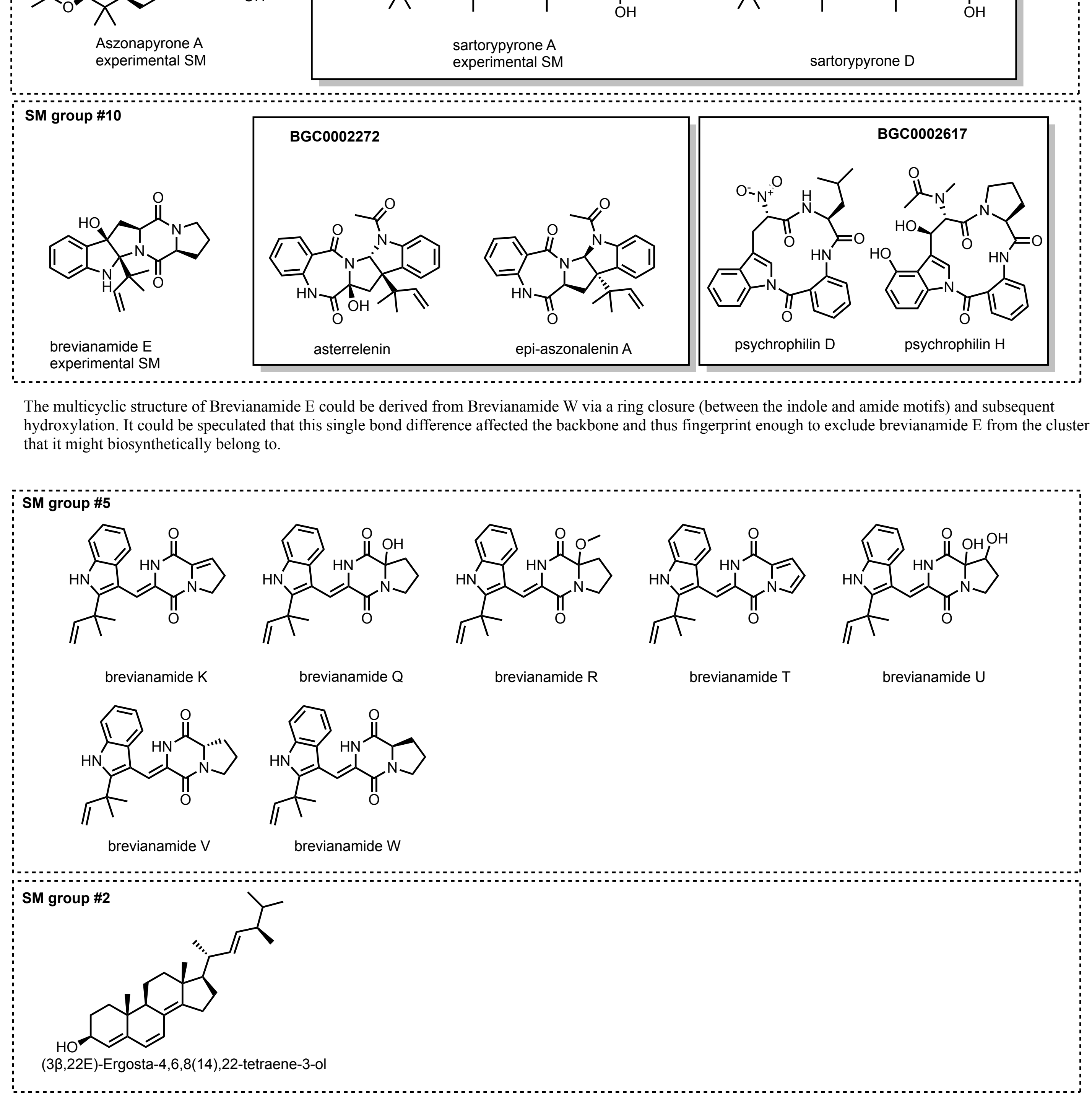

**Figure S5**: Structures of all experimental SM features shown in the assigned SM groups together with similar SM from known BGCs.

#### Supplementary tables: Supplementary_tables.xlsx

Collection of all tables mentioned in the main publication, mainly pertaining to results:

- **Table 1** (as in the main publication)
- **Table S1**: extended Table 1
- **Table S2**: Highest Tanimoto similarity score for each experimental SM with a MIBiG SM (chiral and achiral search)
- **Table S3** BGCs in *A. fischeri*: Evaluation of all names antiSMASH BGC predictions plus the ones found via sequence similarity search with MIBiG amino acid queries.
- **Table S4** BGC information: extensive information on all BGC genes across all strains including genomic location, blast identities to MIBiG queries, classification etc.

#### Raw and intermediate data provided separately

- BGC_grouping_aS7_PROCESSED.csv: Manually corrected results of clustering antiSMASH BGC regions by gene content.
- BGC_grouping_aS7_unprocessed.csv: Initial results of clustering antiSMASH BGC regions by gene content.
- annot.anotations_for_BGCs.extended.txt
- anti.antiSMASH7_unscaffolded.prosmash_output.tsv: results after combining antiSMASH results using in-house script prosmash.py.
- mibig_to_DTO.AA.tsv: Combined results of diamond blast searches of MIBiG v.3.1 amino acid queries to the *A. fischeri* genomes.
- pan.Fischeri_pangenome_info.csv: Pangenome information, i.e. correspondence of genes/transcripts across the 16 strains. Orthologous genes are tracked by the same IDX.

- strain_names.csv: Throughout the project and before receiving official strain designations, the different collaborating labs used internal IDs which are provided here.
- mibig_BGCs_genes.tsv: List of all genes per BGC in MIBiG v3.1
- compounds_for_clustering.csv: SMILES and names of all compounds in MIBiG v3.1 and at the bottom, the experimental SMs found in *A. fischeri*.
- compounds_strain_temperature_count.csv: count of the number of replicates (0–3) in which a given SM was detected in a given strain at a given temperature.

#### Custom analysis code and in-house scripts

- Based on antiSMASH results, we ran in-house bash and python scripts to collect antiSMASH results (run_prosmash.sh, prosmash.py).
- To identifying the same antiSMASH predicted BGCs across the 16 genomes, we compared the BGC regions by gene content using antiSMASH7_BGC_clustering.R.
- For the manual BGC validations, we compiled a markdown detailing all results and decisions (Supplement_BGCs.Rmd)
- Manual script detailing how we clustered compounds using rdkit as documented in compound_clustering_A_fischeri.ipnb
- GUI to find the most similar compound in MIBiG v3.1 for a compound of your choice ()
